# Enzymatically-active bacterial microcompartments follow substrate gradients and are protected from aggregation in a cell-free system

**DOI:** 10.1101/2022.05.16.492142

**Authors:** Jan Steinkühler, Charlotte H. Abrahamson, Jaime Agudo-Canalejo, Ramin Golestanian, Danielle Tullman-Ercek, Neha P. Kamat

**Affiliations:** Department of Biomedical Engineering, Northwestern University, Evanston, IL 60657; Department of Chemical and Biological Engineering, Northwestern University, 9 Evanston, Illinois, USA; Department of Living Matter Physics, Max Planck Institute for Dynamics and Self-Organization, D-37077 Göttingen, Germany; Rudolf Peierls Centre for Theoretical Physics, University of Oxford, Oxford OX1 3PU, United Kingdom; Center for Synthetic Biology, Northwestern University, Evanston, IL 60657; Chemistry of Life Processes, Evanston, IL 60657

## Abstract

The ability to dynamically control organelle movement and position is essential for cellular function. Yet the underlying mechanisms driving this organization have not been fully resolved. Here, we draw from recent experimental observations and theoretical models of enzyme chemotaxis to demonstrate the chemotaxis of a bacterial organelle, the 1,2 propanediol (1,2-PD) utilization bacterial microcompartment (MCP) from *Salmonella enterica*. Upon encapsulating MCPs in a cell-like, biomimetic compartment, we observed the directed movement of MCPs along an external gradient of substrate. Our analysis shows that MCPs not only chemotax towards their substrate but also that enzymatic activity and substrate turnover protect them against large-scale aggregation. Our results provide a first experimental demonstration of organelle chemotaxis in a synthetic cellular system and support a recent theoretical model of chemotaxis. Together this work reveals a potentially significant driver of organelle organization while contributing to the construction of synthetic cell-like materials.

## Introduction

The spatial organization of sub-cellular components is dynamic, requiring rearrangements as cells grow, move, and divide. Yet even between these larger-scale events, organelles and biological molecules move within cells, localizing at specific points in cells to promote synthesis and mediate transport.^1–4^ Recently, a range of enzymes have demonstrated the capacity to move towards or away from substrates.^5–8^ A fundamental question that has emerged is whether the directed motion of catalytically active systems can be harnessed to drive dynamic reorganization of cellular mimetic “organelles”. Such a capability would enable new technologies ranging from targeted drug delivery to self-organizing metabolic networks.^9,10^ Additionally, these effects are expected to be crucial for the assembly of intelligent machines such as synthetic cells.^11^

Interactions between catalytic particles and substrates can result in either positive or negative chemotaxis, towards or away from a substrate.^5,7,12^ This directed motility can be conferred to larger structures, such as lipid and polymer vesicles, when enzymes are encapsulated or localized to them.^10,13^ In previous studies, chemotactic behavior and enzyme activity were coupled, meaning that depending on the choice of enzyme and substrate concentration, chemotaxis up or down the concentration gradient would occur. It would be desirable to decouple these effects, and to obtain a mechanism by which any compartmentalized enzymatic reaction can be targeted towards its substrate.

As a model compartment, we chose the 1,2-propanediol utilization bacterial microcompartment (MCP) from Salmonella enterica serovar Typhimurium, which contains the enzymes required to metabolize 1,2-PD. MCPs are metabolic organelles, 40 to 200 nm in diameter, with a protein shell that sequesters segments of enzymatic pathways.^14^ These shell proteins permit the transit of the substrate through pores to engage in enzymatic reactions in the MCP interior. The residues lining the pores can be mutated to alter enzyme kinetics of encapsulated reactions.^15–17^ MCP enzyme contents can be readily modified to vary catalytic activity and to include additional cargo like fluorescent proteins.^18–20^ MCPs are therefore an ideal platform to investigate the chemotaxis of nanoscale compartments where shell structure, enzymatic activity, and fluorescent contents can be independently tuned.

We first present a coarse-grained computer model that we used to identify important parameters to confer robust compartment chemotaxis, and that was guided by a recently proposed mechanism known as stabilitaxis.^21^ Through comparison to our computational model, we show how catalytic activity protects MCPs against large-scale aggregation, an emergent, autoregulating phenomenon. In this way, we introduce a new mechanism by which chemotactic matter can be protected from aggregation and establish a chemotactic platform that will benefit new biotechnological applications.

## Results

### Computational study of chemotactic compartments

To understand parameters that might influence compartment chemotaxis in our system, we first set up computational studies to monitor the trajectories of spherical compartments as a function of substrate concentration, substrate binding affinity, and substrate consumption rate. Spherical compartments of radius *R* were placed next to smaller particles of radius 0.1 *R* that represented substrate (gray and red spheres in Fig 1A, respectively), with a compartment-substrate binding energy *ε*. Binding of the substrates to the compartment surface induced reversible binding between compartments via compartment-ligand-compartment complex formation, a process we call “crosslinking” (Fig 1B, Fig S1A). Next, we studied a particle-based, reaction-diffusion simulation of *N*=15 compartments in a rectangular simulation box of size 5×5×50 *R* (reflecting boundary conditions) (Fig 1C). Substrate particles were introduced in a stochastic manner at an average rate *k_s_* at one side of the simulation box (red line in Fig 1C, Fig. S1B). The observed trajectory of compartments depended strongly on the parameter ε: when only repulsive interactions were included (*ε* = 0 kcal/mol), the compartments diffused within the rectangular box with no preference for either direction along the long axis of the simulation box. However, if compartments were able to bind to the substrate (*ε* = 10 kcal/mol), a directed movement towards the substrate source was observed. This directed movement resulted in accumulation of the compartments towards the source of the substrate (Fig 1C,D). The directed movement was due to the slower diffusion of larger, crosslinked assemblies of multiple compartments relative to the faster diffusion of free, individual compartments. The differential diffusivities of free and crosslinked compartments led to an accumulation of compartments at the substrate source over time, where crosslinking is favored, consistent with a recently proposed theoretical model.^18^

**Figure 1.**
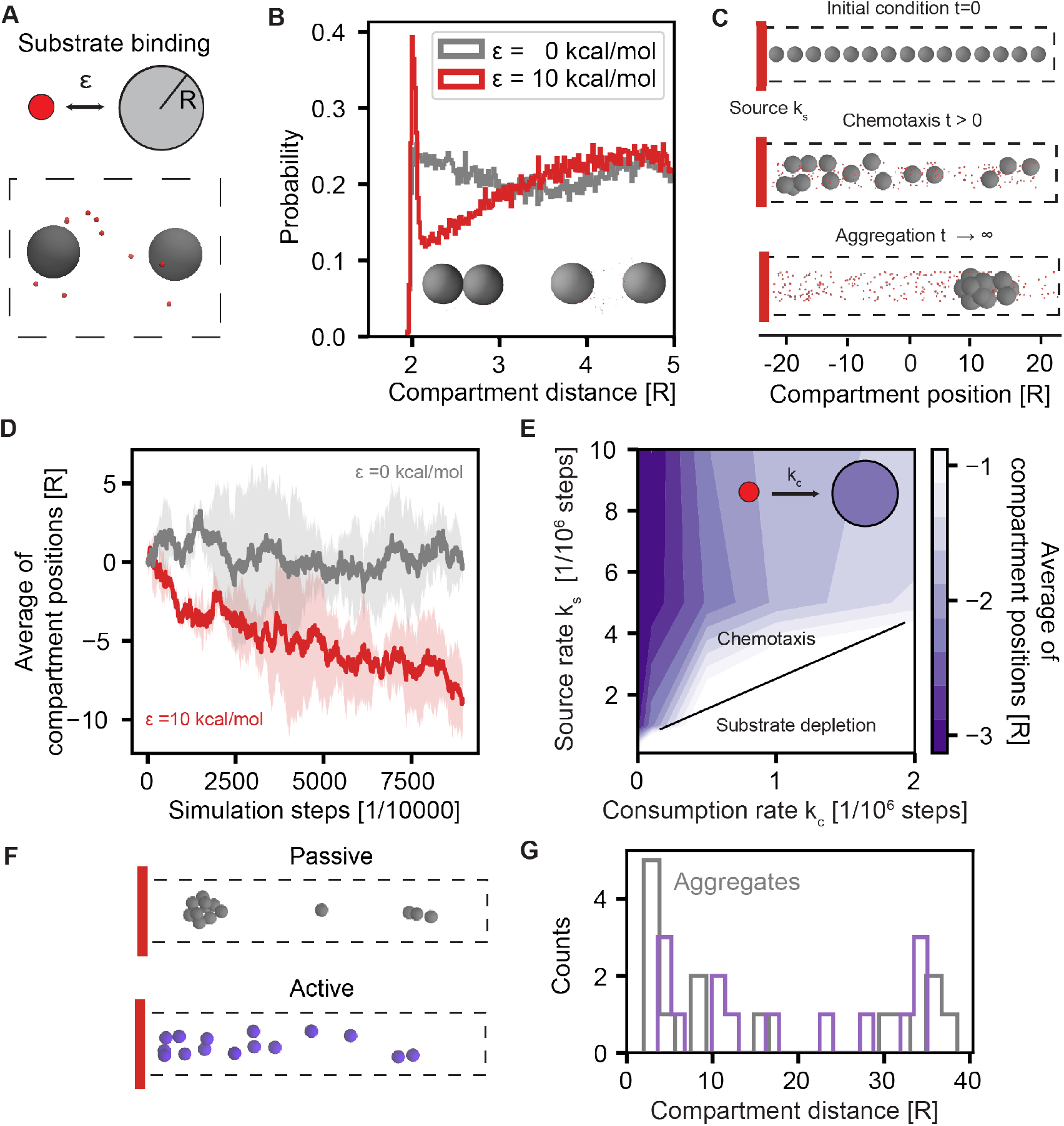
Coarse-grained simulations of chemotactic compartments A) Schematic of components in the simulation which include small substrate molecules (red spheres) that bind to larger compartments of radius *R* (gray spheres) with a binding energy *ε*. The dashed line indicates a simulation box containing two compartments and multiple substrate particles. B) Histogram of the distance between compartments when binding energy between compartment and substrate particles is ε = 0 (gray) or ε = 10 kcal/mol (red). C) Example snapshots of simulations in which multiple compartments are placed inside a box with reflective boundary conditions (dashed gray lines) and a substrate source (red) for ε = 10 kcal/mol. (Top) Initial configuration of compartments, (Middle) compartments demonstrate chemotaxis at intermediate time points, and (Bottom) compartments eventually aggregate at long time intervals. Coordinate system parallel to the long side of the box is shown units of the compartment radius *R* D) Example trajectories for the average compartment positions along the long side of the simulation box (see panel C for coordinate system) for two values of substrate interaction energy, ε. The average positions were calculated by the geometric mean over all compartments. Shaded region indicates std. dev. of four replicates. E) Color map indicates the average compartment positions at the end of the simulation as a function of the rate of substrate introduction (*k_s_*) and consumption (*k_c_*). (In this panel *ε* was 50 kcal/mol to improve sampling at low particle concentrations). The map predicts conditions of substrate introduction and consumption that lead to compartment chemotaxis shown by negative values of compartment average positions, indicating enrichment at the substrate source, see panel C and D F) Example simulation snapshots showing aggregation of passive compartments (gray) that do not consume substrate, *k_c_* = 0, and active compartments (violet) that do consume substrate, other parameters were *k_c_* = 0.5·10^-6^/steps, *ε* = 50 kcal/mol, and *k_s_* = 5·10^-6^/steps. G) Corresponding comparison of compartment spatial distribution as a function of substrate consumption rate. Histogram shows compartment location for data collected after 4·10^9^ integration steps, n=3 replicates. In all panels, distances and positions are shown in units of compartment radius *R*, see panel C for definition of the coordinate system.

The chemotactic behavior of particles in our simulation had a finite lifetime. We observed the directed motion of compartments stopped upon compartment aggregation. Over time, as more and more substrate particles were introduced in the simulation box, the crosslinking effects became stronger, and eventually all compartments were aggregated in a large cluster (Fig 1C). These large aggregates did not accumulate at the particle source because their association was no longer dependent on the gradient of the substrate. Aggregation, therefore, appears to be a limiting factor for sustained chemotactic behavior.

We wondered what effect catalytic activity would have on compartment movement and aggregation. To represent this phenomenon, we modelled “active” compartments in which substrate molecules that were bound to the compartment surface were removed with a rate *k_c_*. In contrast, “passive” compartments were those with no substrate removal (*k_c_* = 0). In a general sense, substrate uptake might represent passive uptake through compartment pores followed by enzymatic turnover into products that do not further interact with the substrate and compartments. This consumption term allowed us to study the evolution of the system as a function of the two rates, *k_s_* and *k_c_*, where we color-coded the average center of mass of the compartments across the simulation trajectory (Fig 1E). As expected, for very small rates of substrate activity, *k_c_* ≪ *k_s_*, the chemotactic behavior and subsequent aggregation at high substrate concentrations was observed. However, in the case of fast substrate uptake, *k_c_* > *k_s_*, no substrate gradient developed, and compartments were, on average, uniformly distributed in the simulation box (region of substrate depletion, Fig 1E). An interesting case was observed for intermediate uptake rates where the total uptake rate by all *N* compartments was comparable to the rate at which substrate appears, *Nk_c_ ~ k_s_*. For these cases, the number of substrate particles seemed to converge to a steady state value (Fig. S1C). This autoregulating behavior reflects a balance between chemotaxis toward the substrate source and locally enhanced substrate consumption, which reduces crosslinking and counteracts chemotaxis.

Importantly, sufficiently active compartments still underwent chemotaxis towards the source but now avoided large-scale aggregation. The capacity of substrate depletion to counteract compartment aggregation can be seen by considering the distances between compartments at the end of the simulation trajectories. Most of the compartments were found within a cluster for passive compartments (*k_c_* = 0), but were found more loosely packed for active compartments (Fig 1F,G). Taken together, our simulations indicate that a balance between binding affinity to a substrate and consumption of that substrate is needed to sustain chemotactic behavior of compartments and avoid large-scale aggregation.

### Experimental realization using bacterial microcompartment

We next set out to experimentally realize the directed movement of chemically active compartments. We used a recently introduced genomic expression platform to obtain two types of purified MCPs from Salmonella enterica serovar Typhimurium.^18^ These two types of MCP were “active” or “passive” depending on their capacity to consume a 1,2-PD substrate. Active MCPs included the enzymes diol dehydratase (PduCDE) and 1-propanol dehydrogenase (PduQ) (Fig 2A, full name ΔpduP::ssPGFPmut2, later called “active”), which yields the product 1-propanol. Passive MCPs did not include PduCDE and thus were not active on the substrate 1,2-PD (ΔpduD::ssDGFPmut2, later called “passive”). Both types of MCPs were engineered to carry either fluorescent GFP or mCherry proteins in high copy numbers by using signal sequences from natively encapsulated Pdu enzymes. Importantly, both types of MCPs consist of the same protein shell, providing a platform in which enzymatic function and fluorescent visualization can be tuned independently while maintaining compartment structure and size. The active MCPs consume 1,2-PD and the passive MCPs do not consume any 1,2-PD. To pursue our computational predictions, we set out to observe the potential for *in vitro* reconstituted MCPs to chemotax towards the 1,2-PD substrate as a function of their enzymatic activity.

**Figure 2.**
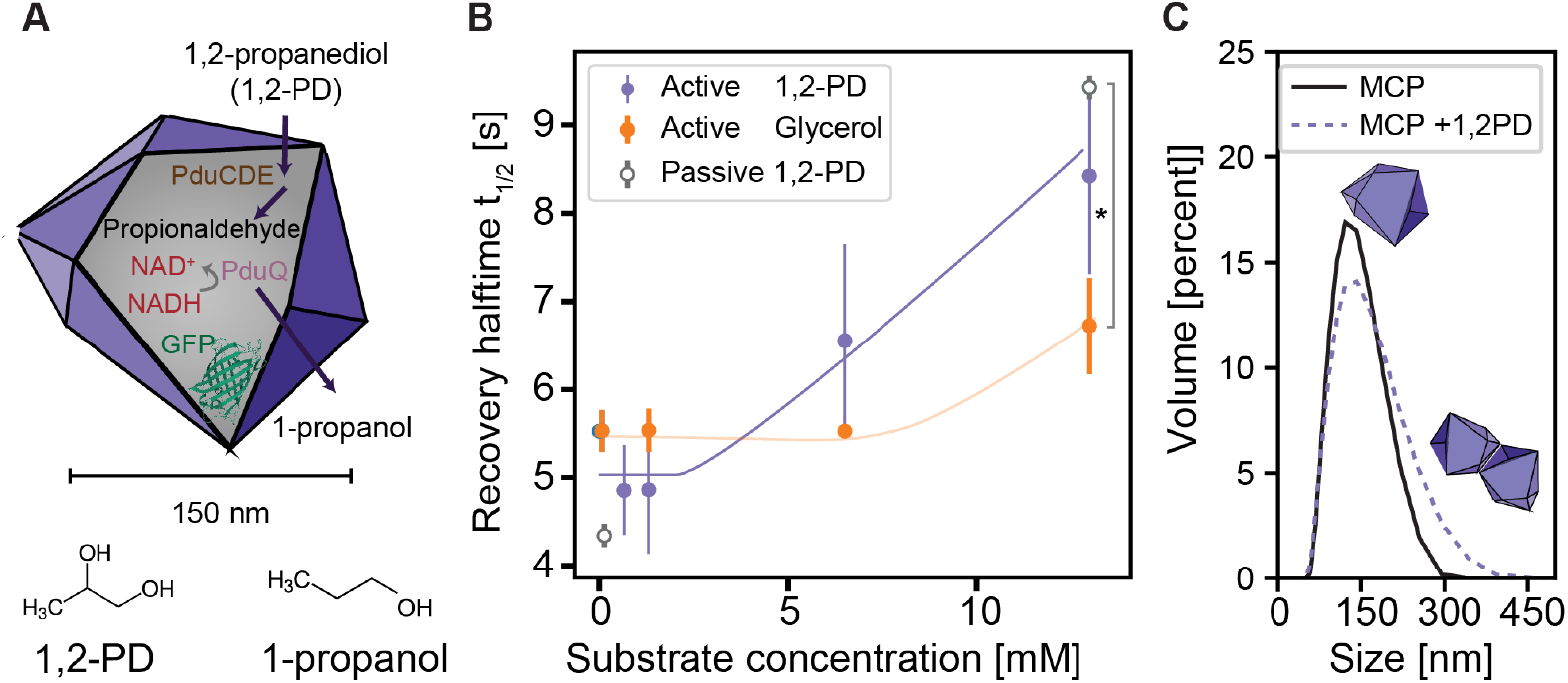
Substrates induce crosslinking of both active and passive compartments. A) Schematic of the reaction pathway of an active Pdu bacterial microcompartment (MCP). B) Fluorescence recovery after photobleaching (FRAP) measurements of fluorescent MCPs as a function of substrate concentration for MCPs with or without activity. Recovery times (inverse of the diffusion constant) show both active and passive compartments bind substrate in a concentration dependent manner. At zero substrate concentration the measured recovery times correspond to a diffusion constant in the order of 5 μm^2^/s. Changing the substrate to glycerol requires more substrate to slow MCP diffusion to a similar extent as the native 1,2-PD substrate. Error bars indicate std. dev. for n=3 repeats, * p < 0.05, p values were calculated using a Student’s t-test. C) Light scattering measurements of active MCPs in the absence or presence of 1,2-PD (1.3 mM) show a shift to larger sized particles in the presence of 1,2-PD, suggesting MCP-MCP crosslinking. (Histogram obtained from n=3 repeats).

### MCP are crosslinked by their substrate in a concentration-dependent manner

We first investigated the effect of substrate identity and MCP activity on MCP crosslinking when MCPs were free in solution. Purified MCPs were mixed with substrate (1,2-PD) at room temperature and examined under the microscope. Using photobleaching experiments, we measured the fluorescence recovery time, t_1/2_, after exposing a fraction of the solution volume to a strong laser pulse. The half time provides a measure of the time that is needed for diffusional exchange of irreversibly bleached (dark) MCP with the pool of fluorescent MCP surrounding the bleached volume (Fig S2A). With 1,2-PD, the natural substrate for the active MCP, increasing substrate concentration led to longer recovery times, indicating slower diffusion of MCPs (violet Fig 2B). Because only low millimolar concentration of substrate were used, we could rule out any viscosity or solvent effects as the cause of this slowdown. As in our computational model, we attribute the slower diffusion to the crosslinking effect of passive MCPs (gray points Fig 2B), suggesting that it is the compartment shells that interact to form crosslinked complexes. Pdu MCP shells consist of nine different proteins, with the proteins PduA, B, B’, and J as the major shell components.^14^ In principle, any of these proteins might exhibit weak multivalent interactions with the substrate to induce crosslinking. Again, we emphasize that all shell proteins are present in both active and passive MCPs which should lead to similar binding affinities to 1,2-PD. We further studied glycerol (orange Fig 2B), which, like 1,2-PD, permeates MCPs via PduA.^16^ Interestingly the glycerol MCP crosslinking efficiency was significantly lower (* in Fig 2B) compared to 1,2-PD and not observed for intermediate glycerol concentrations.

These results can be used to discuss the relation between the coarse-grained simulation and our experiments. We focus our discussion on PduA, which assembles as a homohexamer with a central pore and is thought to mediate selective passage of 1,2-PD into the MCP.^19,20^ We first recap what is known for the molecular mechanism of substrate passage via PduA. Selectivity of PduA towards the substrate is likely to be mediated by the residues lining the central PduA pore.^16,22^ Previous studies estimated that the PduA pore provides a 1,2-PD binding energy of about 1 kcal/mol mainly mediated via hydrogen bonding to the PduA residue S40 at the entry of the pore.^23^ The surface bound state then facilitates substrate translocation via diffusion through the pore. In addition to substrate translocation, a transiently, surface-bound substrate provides the opportunity to induce a crosslinking effect between multiple MCPs. With an estimated surface density of one hundred PduA proteins per single facet on a MCP^14^, the maximum expected binding energy is well within the computationally-studied values of ε. Compared to 1,2-PD, less favorable interactions between glycerol at the entry of the pore lead to less efficient glycerol influx.^16^ Indeed, reducing ε diminishes cross-linking efficiency in our model, consistent with the experimental condition of using glycerol as a substrate. This comparison shows that our coarse-grained parameter ε provides a sufficient description of the experimentally observed MCP crosslinking effect.

To further confirm MCP crosslinking, we used a second experimental method of dynamic light scattering which does not rely on fluorescence output but measures the correlation time observed by light scattering of MCPs. Consistent with our bleaching experiments, we observed a fraction of MCP dimers after incubation with 1,2-PD (Fig 2C). Together, our observations indicate that crosslinking of MCPs is mechanistically independent of the chemical activity of the encapsulated enzymes and instead is determined by the crosslinker concentration and its ability to bind the shell proteins, fulfilling an important design goal to decouple chemotactic directionality and enzymatic activity.

### High-throughput encapsulation of MCPs in liquid-liquid phase separated droplets

We next set out to investigate the capacity of a substrate gradient to induce the directed movement of MCPs. In our simulations, a substrate gradient was simply generated by introducing particles at one side of the simulation box. We hypothesized that a similar setup would allow us to study MCP chemotaxis experimentally. To expose MCPs to a gradient of substrate concentrations, we used liquid-liquid phase separation. Briefly, when a substrate is soluble in both phases but is initially present in only one of the phases, it will diffuse over time into the other phase via diffusion across the interface between the phases. Accordingly, we hypothesized that by placing an organic 1-octanol phase containing 1,2-PD in contact with an aqueous phase containing MCPs, we could create conditions that will lead to a gradual increase of 1,2-PD in the aqueous phase. Indeed, this effect was observed when a macroscopic liquid droplet of 1-octanol carrying 1,2-PD was placed in contact with an aqueous phase carrying MCPs. Over time we observed large clusters of crosslinked MCPs that could be directly visualized using confocal microscopy (Fig. S2B), indicating 1,2-PD was able to diffuse from the 1-octanol phase into the aqueous phase. Control experiments lacking 1,2-PD did not lead to such cluster formation, confirming that crosslinking of MCPs into clusters was dependent on the presence of substrate. These experiments demonstrate that liquid-liquid phase separation is in principle a suitable route to introduce a substrate gradient in the MCP environment. However, we found that experimentally generating a sustained, linear gradient in macroscopic liquid droplets was challenging because the preparation and handling would lead to perturbation of the diffusive front. We therefore sought to assemble cell-sized compartments that encapsulate MCPs in a liquidliquid phase-separated system that would better maintain and protect a linear gradient of substrate.

To reach this goal, we utilized a flow-focusing microfluidic system to generate monodisperse, cell-sized, water droplets containing MCPs in a continuous organic phase of 1-octanol (Fig 3A). To stabilize the water-oil interface we included a variety of amphiphilic molecules in the 1-octanol phase. These amphiphiles self-assembled into a monolayer at the water-oil interface which provided a biomimetic, stabilized interface. The amphiphilic monolayer further served to prevent adsorption of MCPs to the water-oil interface (Fig S3A). Phospholipids, which outcompete the MCPs for the interfacial location, enabled the free diffusion of MCPs in the droplet lumen as shown in Fig. 3A and Fig. S3B. We also observed that other macromolecules could be used in place of phospholipids to stabilize the water droplet interface and avoid nonspecific MCP adsorption, including Poloxamer 188, a synthetic tri-block copolymer consisting of hydrophobic/hydrophilic blocks (Fig S3C). Other more hydrophilic polymers like 8 kDa polyethylene glycol did not stabilize MCPs away from the liquid/liquid interface (Fig S3D). Apart from providing droplet and MCP stabilization, we observed that the tri-block copolymer is beneficial by acting as a crowding agent, reminiscent of the crowded cytosolic environment, leading to slower and experimentally better accessible diffusion timescales. We anticipated this cellular mimetic compartment encapsulating MCPs and stabilized with a crowding polymer at the interface would allow us to study MCP chemotaxis, and therefore used the Poloxamer 188 molecule to stabilize compartment interfaces for subsequent studies.

**Figure 3.**
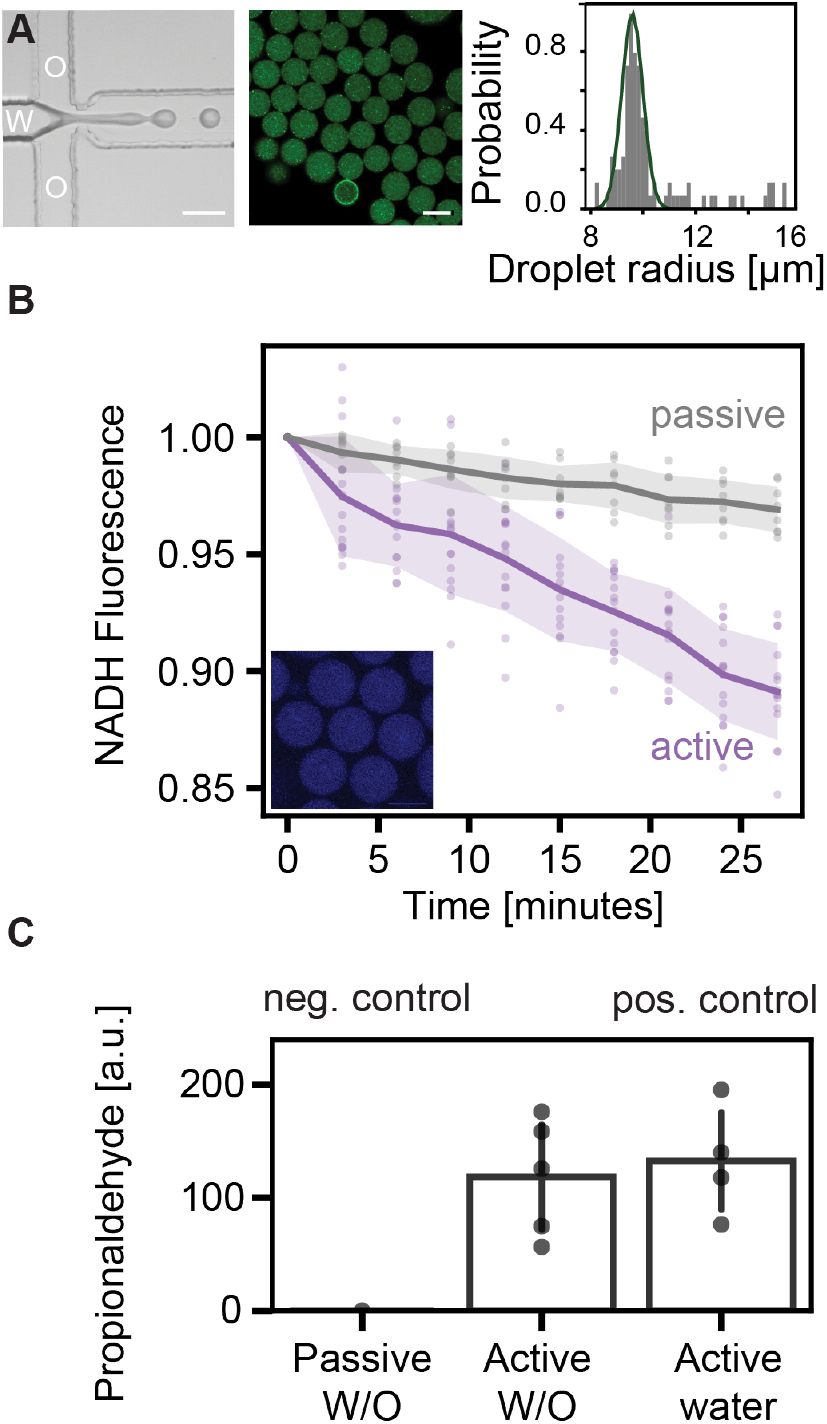
Microfluidic encapsulation of MCPs into cell-sized compartments enables spatial assessment of MCP activity. A) (Left) Phase contrast microscopy image of a microfluidic device used to assemble MCP-encapsulated compartments. A water phase (W) containing MCPs breaks up into droplets within the continuous 1-octanol phase (O). (Middle) Representative fluorescence microscopic image of compartments containing fluorescent MCPs (green) with a corresponding droplet size distribution (Right). B) NADH fluorescence from individual droplets over time when 1,2-PD is present in the oil phase for active (purple) and passive (gray) MCPs. (Inset) Droplets with NADH fluorescence. C) Detection of reaction intermediate propionaldehyde in the oil phase when passive MCPs or active MCPs were encapsulated in water droplets alongside a noemulsion positive control in which MCPs are directly mixed with 1,2-PD in a water phase. Each datapoint from a repeat experiment, error bar indicates std. dev. Scale bar is 20μm

We next confirmed that the substrate 1,2-PD could indeed cross the droplet interface to interact with MCPs when 1,2-PD was introduced in the continuous octanol phase. To assess turnover of 1,2-PD by active MCPs, we monitored the conversion of NADH, intrinsically fluorescent in visible wavelengths, to non-fluorescent NAD^+^ by the NADH-dependent dehydrogenase reaction that converts the intermediate propionaldehyde to the product 1-propanol (PduQ in Fig 2A). 1,2-PD was added to the 1-octanol phase and NADH fluorescence, which should decrease upon 1,2-PD consumption, was measured. We observed that NADH levels substantially decreased in the presence of active MCPs relative to passive MCPs (Fig 3B). This result suggests that MCPs remain catalytically active throughout our microfluidic assembly procedure and that 1,2-PD added to the outside of the water droplets can cross the droplet interface to access MCPs. We further confirmed the activity of droplet-encapsulated MCPs via high performance liquid chromatography (HPLC). Interestingly, for active MCPs we observed the exchange of propionaldehyde from the MCP interior to the octanol phase. As expected, passive compartments did not generate detectable propionaldehyde levels. We compared the level of propionaldehyde in the oil phase to a noemulsion control of two macroscopic phases of MCP in water in contact with 1-octanol, finding similar propionaldehyde concentrations (Fig 3C). These results show that MCP activity was preserved during the encapsulation procedure as well as the ability of the phase separated system to scavenge reaction intermediates from the MCP interior. Altogether our system allowed for controlled substrate addition to stable and chemically active MCPs, allowing us to experimentally investigate the effect of a substrate gradient on MCP movement.

### Compartmentalization in cell-like droplets enables MCPs to chemotax towards external gradients of MCP substrate

To examine the chemotactic movement of MCPs in response to a substrate gradient, we added two aqueous droplets to a continuous 1-octanol phase. One droplet contained the 1,2-PD substrate and a tracer dye to enable tracing of the 1,2-PD location over time. The second droplet contained MCPs. Specifically, we pipetted a relatively large 500 nL aqueous droplet with 131 mM 1,2-PD into the continuous 1-octanol phase acting as source of 1,2-PD. The source droplet contained a fluorescent dye that is soluble in 1-octanol and serves as an indicator for 1,2-PD location inside the 1-octanol phase (cyan in Fig 4A). The fluorescent dye is about 8 times larger in molecular weight compared to 1,2-PD, making the dye signal a qualitative indicator for the direction of the 1,2-PD diffusive front. We then used confocal microscopy to visualize the resulting diffusion of the tracer dye and MCP distribution. The time needed for the diffusive front to reach each MCP droplet varied as a function of the distance between the source and target droplet, and was generally between 5 and 30 minutes. Due to the relatively slow diffusion in 1-octanol, each analyzed droplet could be selected to have developed a similar 1,2-PD gradient as indicated by the tracer dye.

**Figure 4.**
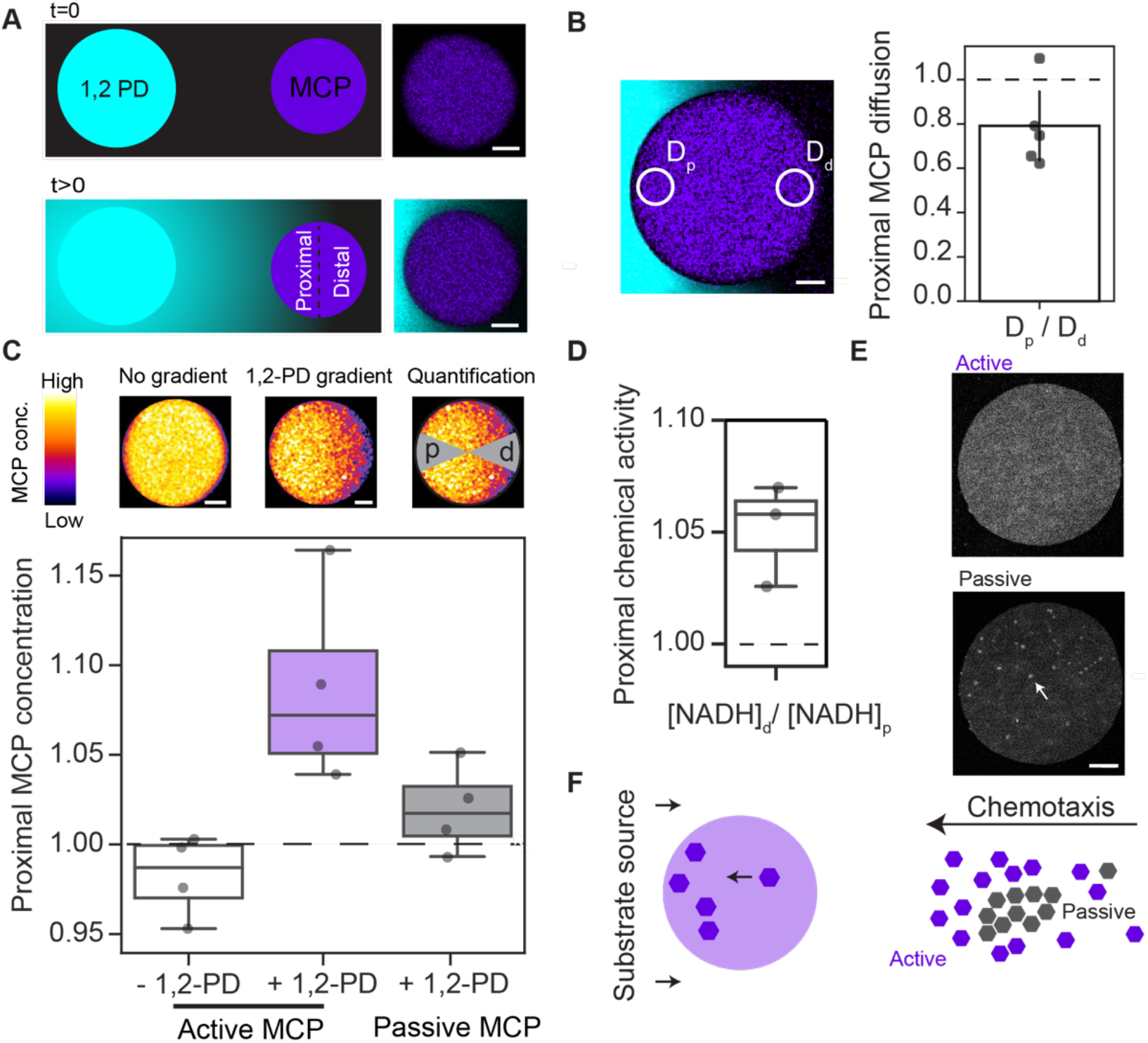
MCPs chemotax along a gradient of 1,2-PD. A) Cartoon showing experimental setup with source water droplet containing 1,2-PD and tracer dye (Atto 647N, shown in cyan) and target MCP-containing droplet (violet). Black is the continuous 1-octanol phase. Over time, a diffusive gradient of 1,2-PD and tracer dye propagates through the 1-octanol phase. Fluorescent microscopy images from experiment are shown adjacent to the cartoon, scale bar is 20 μm. B) FRAP measurements of the proximal (p) and distal (d) sides of a droplet containing fluorescent, active MCPs. Bar plot shows data from individual water droplets and is reported as the ratio of diffusion constants at the proximal to distal regions of the droplet. Ratios of fluorescence recovery times below one indicates MCPs on the proximal side diffuse slower than the distal side (Student’s t-test p<0.05). Std.dev. indicated by error bar, n=5 repeat experiments. Scale bar 10 μm. C) Contrast-enhanced MCP fluorescence (GFP fluorophore) shown in ‘hot’ color map in conditions of absent and developed 1,2-PD gradient. Scale bar is 10 μm. Fluorescence intensity measurements of MCPs in water droplets are reported as a ratio at the proximal to distal regions (indicated by gray cones) with gradient developed across MCP droplet as indicated by tracer dye. Active and passive compartments are preferentially localized at the proximal region in the presence of a 1,2-PD gradient and are uniformly distributed in the absence of 1,2-PD (Student’s t-test p < 0.05) D) Quantification of NADH fluorescence in droplets containing active MCPs at the proximal to distal locations indicates lower NADH concentration at the proximal droplet side in the presence of 1,2-PD, which in turn indicates higher chemical activity at the proximal end. The chemical activity was taken to be inversely proportional to NADH concentration and thus the ratio between NADH concentration distal to proximal in shown. All datapoints in panels C,D are individual measurements of droplets from a minimum of n=3 repeat experiments. Boxes show 25th and 75th percentiles. E) Fluorescent microscopy images of the same droplet containing both passive MCPs (mCherry fluorophore) and active MCPs (GFP fluorophore) in the presence of 1,2-PD. Aggregates are present when passive MCPs are encapsulated in contrast to active MCPs which appear homogenous. Scale bar is 40 μm. F) Schematic of chemotaxis and aggregation behavior.

We first observed that MCP diffusion became spatially inhomogeneous within the droplet, consistent with 1,2-PD-mediated crosslinking of MCPs. We measured the recovery time after fluorescent bleaching and found that MCPs on the 1,2-PD facing side (proximal) of the water droplets exhibited a lower diffusion constant than those on the farthest side from the 1,2-PD droplet (distal) (Fig 4B), consistent with the bulk measurements at varying substrate concentrations (see Fig 2B). As predicted in our simulations, this situation leads to polarization of the droplet, where MCPs are enriched or ‘focused’ at the proximal side (Fig 4C). We would expect that, in the presence of a substrate gradient, MCP biochemical activity likewise exhibits a focused gradient, increasing net substrate turnover closer to the substrate. Indeed, NADH concentration is reduced at the proximal side of water droplets, reflecting enhanced enzymatic activity where 1,2-PD concentrations are the highest (Fig. 4D and SI Fig. S4A). Consistent with our simulation, active MCPs indeed follow substrate gradients leading to enhanced chemical activity towards the substrate source.

We hypothesized that a recently proposed theoretical model known as stabilitaxis might be used to understand the chemotaxis of compartmentalized enzymatic reactions. Stabilitaxis proposes that an oligomeric protein will drift towards regions in which proteinprotein interactions are favored.^21^ Yet, to date, this model has not been experimentally validated. The measured relative differences between the diffusion coefficient of active MCPs in the proximal and distal sides can be related to the relative accumulation on each side through the general theory of stabilitaxis.^21^ Indeed, in the limit of weak gradients and weak MCP-MCP binding, as expected for active MCPs which self-regulate the local concentration of 1,2-PD crosslinkers, we find the relation 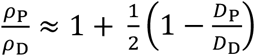 where *ρ*_P,D_ and *D*_P,D_ are the MCP concentrations and effective diffusion coefficients in the proximal and distal sides, respectively (see SI for details of the calculation). For the measured value 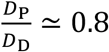 (Fig. 4B), we thus predict that 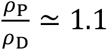, which coincides with the experimental result (Fig. 4C) without any adjustable parameters.

We next examined in detail how the activity of MCPs influenced their chemotactic behavior. We found that both active and passive MCPs follow the substrate gradient, but exact comparison of the magnitude of the effect was difficult at this stage as in our experimental setup each droplet is exposed to slightly different 1,2-PD concentrations. We addressed this limitation by including two MCP populations within each droplet that contained spectrally distinct fluorescent proteins. As before active MCPs contained green fluorescent protein (GFP), but passive ones contained the fluorescent protein mCherry (ΔpduD::ssDmCherry), allowing us to differentiate active and passive MCPs in the same droplet. We confirmed that, within the same droplet, both active and passive MCP show chemotactic behavior of comparable magnitude (Fig. S4B). However, large differences between the compartments were observed in the long-term behavior at higher 1,2-PD concentration. Here, red, bright fluorescent puncta indicated aggregation of passive MCP (arrows in Fig. 4E). As predicted, active MCPs in the same droplet did not aggregate under these conditions (bottom micrograph in Fig. 4E). These results agree with our computational prediction that substrate turnover and consumption are important to mitigate and prevent large-scale aggregation, ultimately sustaining a spatial gradient of chemotaxing MCPs and an associated gradient of chemical activity (Fig. 4F).

## Conclusion

In this work we have studied chemotaxis of organelles using computer simulations, theory, and experiments. Together, these studies provide compelling evidence that the bacterial organelle, the 1,2 propanediol (1,2-PD) utilization bacterial microcompartment, chemotaxes when removed from the cellular environment and is placed in a gradient of 1,2-PD substrate. This chemotaxis occurs via a stabilitaxis mechanism arising from substrate-induced crosslinking of compartments that slows diffusion of the complex relative to individual compartments and leads to compartment localization in regions of higher 1,2-PD concentrations. Chemical activity and the ability to turn over 1,2-PD, in turn, protects against compartment aggregation.

Stabilitaxis is based on the substrate-dependent stability of slowly diffusing complexes made from crosslinked compartments. The result is a net movement of particles towards areas with higher concentrations of crosslinking substrate. This phenomenon might drive the directed movement of a range of different systems, not only of catalytically active biomolecules and organelles, but also of passive-binding particles including beads or lipid vesicles. Additional complexity was introduced in our study by the ability of the compartments to take up and consume the crosslinking molecule. Again, this capacity to reverse compartment crosslinking might be provided by a range of different molecular mechanisms, including transport of the crosslinking molecule through selective pores, by competitive binding interactions with another molecule, and by enzyme-based catalysis.

Our approach to substrate gradient generation utilized a liquid-liquid phase separated water-in-oil emulsion. Several features of this system proved advantageous: first, by assembling cell-sized compartments that contained MCPs, we were able to expose the MCPs to a sustained, linear gradient of substrate that originated from a nearby source, and in which gradient distortions were limited. Second, by using tri-block surfactants to stabilize the water droplet interface, we not only prevented MCP destabilization at the interface, but also varied the physical environment of the droplet to match the crowding conditions of the cytosol that would affect substrate and compartment diffusion and transport dynamics. Third, by utilizing liquidliquid phase separated systems, we provide a route to better control the exchange of substrates and products between the enzyme-containing water phases and a surrounding octanol phase. As liquid-liquid extraction processes are routinely used in industrial biomanufacturing procedures, our system may similarly allow for improved reaction rates and product extraction by taking advantage of differential solubilities of substrates and products in water and oil, respectively.

Biological gradients, like those of oxygen, proteins, and ions, are a hallmark of biology, constantly forming and dissipating in living cells. Our results support a growing body of work that molecular chemotaxis may be an important underlying factor to consider in living systems, not only for small molecules and enzymes, but also for organelles. In this regard, MCPs serve as a unique and powerful tool in the investigation of organelle movement—their ability to be genetically designed allows decoupling between chemotactic behavior and the enzymatic activity encapsulated in the MCPs. The observed crosslinking of Pdu MCPs at millimolar substrate concentrations is consistent with dissociation constants obtained by isothermal titration calorimetry for substrate binding to the EutL protein, a homolog and evolutionary distant shell protein of PduB that could reflect similar associations with PduA.^25^ This observation with EutL suggests the potential applicability of our results to other microcompartment types. What remains unclear is if the studied effects play a direct role *in vivo*, because cytosolic 1,2-PD gradients might vanish quickly. However, we note that the location of carboxysomes has been shown to be controlled by a Brownian-ratchet mechanism leading to the directed movement of carboxysomes. Here, the *in vivo* movement is based on carboxysome-nucleoid binding, mediated by cytosolic gradients of a small crosslinking protein.^26^ The carboxysome-nucleoid binding capacity might therefore be comparable to the simpler substrate-induced, crosslinking-mediated, directed motion of MCPs shown here.

The directed motion of enzymatically active microcompartments in cell-free settings should advance the design of biotechnologies with self-sorting capabilities, such as artificial organelles and cells. Specifically, our approach is a blueprint to produce synthetic cells that encapsulate modular “organelles” with enzymatic pathways, couple mass transport to chemical activity, and allow for spatiotemporal control over chemical activity via chemotaxis. The end result is the dynamic creation of a polarized synthetic cell, in which molecular constituents and the location of chemical activity has reorganized in response to an externally applied chemical gradient. This capability should provide new solutions for challenges in metabolic engineering, biocatalysis, and bionanotechnology.

## Methods

A detailed description of the methodology can be found in the Supplementary Information. A short description is given here.

### Computer simulations

Particle-based reaction-diffusion simulations were performed using the ReaDDy engine.^27^ Compartments were modelled as large spheres that were able to bind to small spheres (substrate) with binding energy ε. The simulation setup varied between simulations with constant number of substrate particles, substrate added at a constant rate and/or substrate uptake by compartments.

### General materials

Microcompartments were purified using differential centrifugation according to previous methods.^18^ All chemicals were obtained from Sigma.

### Data recordings and analysis

Purified MCPs were in suspended in buffer containing cofactors NADH and Ado-B12. MCPs were studied in bulk or as droplets encapsulated in 1-octanol with Poloxamer 188 as a stabilizing surfactant. Confocal images and FRAP data were obtained on a Nikon Eclipse Ti2 confocal microscope. MCP fluorescent distribution were analyzed using Fiji by two 45° segments of the circular MCP droplet as indicated in Figure 4C.^28^ These and all other methods are described in more detail in the Supplementary Information.

## Acknowledgments

This work was supported in part by the National Science Foundation under Grant No. 1844336 (to (to NPK, DTE, CHA, and JS) and Grant No 2145050 to NPK. This research was supported in part through the computational resources and staff contributions provided for the Quest high performance computing facility at Northwestern University which is jointly supported by the Office of the Provost, the Office for Research, and Northwestern University Information Technology.

