## Supporting Information for "Enzymatically-active bacterial microcompartments follow substrate gradients and are protected from aggregation in a cell-free system"

#### Supplemental Methods

##### Computer simulations

Particle-based reaction-diffusion simulations were performed using the ReaDDy engine. We considered spherical compartment particles that interact with each other via a harmonic repulsion potential, corresponding to a "soft" particle radius $R$ with a force constant of 5 kcal/mol. The small substrate particles interact with the compartment particles via a weak piecewise harmonic potential with a cutoff and an energetic minimum of depth $\varepsilon$. In physical units, the simulations correspond to a particle of radius $R$ = 30 nm with a diffusion constant of 0.01 $nm^2$/ns at a temperature of 293 K, roughly corresponding to the experimentally studied system. Substrate-compartment complexes had a preferred distance of 31 nm with the interaction potential cutoff at 33 nm. Substrate particles had a diffusion constant of 0.1 $nm^2$/ns. The simulations were performed with a 0.1 ns timestep. Boundary conditions were either periodic with a box size 200x200x200 $nm^3$ (Fig. 1B) or confined to a rectangular box of 150x150x1500 $nm^3$ (Fig. 1C and following). For gradient formation, we considered 15 substrate particles. In the following chemotaxis simulations, substrate particles were introduced using a Gillespie reaction scheme. Time integration was performed using isotropic Brownian dynamics.

##### MCP production and isolation

Microcompartments were purified using differential centrifugation according to previous methods(Nichols et al., 2020). Briefly, strains were grown at 30˚C for 24 hours in LB-L media. 200 uL of the 24 hour cultures were then added to 200 mL of NCE media (No Carbon Essential) with 55 mM 1,2-propanediol to induce MCP formation. Cultures were then grown for 18-20 hours, until cultures reached at least OD 1. Resulting cultures were then spun down at 5000 RPM for 5 minutes at 4 C. Resulting pellets were lysed chemically using an octylthioglucoside (OTG) solution. The lysate was clarified at 12,000 x g for 5 minutes at 4 C to remove cell debris. The supernatant was then spun at 21,000 x g for 20 minutes at 4 C to pellet MCPs. Pellets were resuspended in 150 uL buffered saline solution containing 50 mM Tris (pH 8.0), 50 mM potassium chloride, and 5 mM magnesium chloride. The resulting product was then assayed for protein content using a BCA assay (Thermo Fisher) and stored at 4C until needed.

**Bacterial microcompartments (MCPs) reconstitution experiments**

After purification MCPs were stored for up to a week in isolation buffer at 4°C. Before experiments MCP were diluted to a concentration of 200 µg/mL in phosphate buffered saline. If not indicated otherwise, the Poloxamer 188 was added to a concentration of 100 mg/mL. Co-factors were supplemented to yield a final concentration of 3 mM (NADH, Sigma-Aldrich) and 4.5 µM (Ado-B12, Santa Cruz Biotechnology). Before experiments, solutions were stored on ice.

**Fluorescent recovery after photobleaching (FRAP)**

FRAP experiments were performed on a Nikon Ti2 confocal microscope using a 20x (Plan Apo) lens. MCPs were prepared as described above and substrates were added at the indicated concentration before FRAP experiments. Then a 30 µL solution was pipetted on a glass slide and a chamber was formed by a top cover glass and "press-to-seal" silicon spacers (Sigma-Aldrich). After a 10-minute equilibration time, a circle of 25 µm in diameter was bleached using the 488 laser line for 50 consecutive frames and recovery was observed under normal imaging conditions for 150 frames and 2 frames per second.

**Dynamic light scattering (DLS)**

MCPs were prepared in Tris buffer as described above and measured in a micro-cuvette (100µL volume) after 10-minute equilibration time at room temperature. Measurements were performed using MALVEN Zetasizer Nano ZS instrument. Obtained Autocorrelation curves were checked for satisfactory fit to the model and volume distribution was reported.

**Macroscopic interface formation**

1-octanol (Sigma-Aldrich) was pre-equilibrated against Milli-Q water stirring equal volumes of both phases, and equilibrated overnight. Before experiments, 1-octanol was pipetted from the upper phase. This "wet" 1-octanol is referred to as "pre-wetted 1-octanol" further below. As indicated in some experiments, 131 mM 1,2-propendiol (Sigma-Aldrich) was added with the Atto 647N tracer dye. Two 25 µL droplets of either 1-octanol or aqueous MCP solution were pipetted on a glass cover slide. Upon contact formation a chamber was formed by a top cover glass and "press-to-seal" silicon isolators (Sigma-Aldrich). The interface between MCP solution and 1-octanol was then imaged over time.

**Microfluidic droplet formation**

Microfluidic chips, model F01-RAW, were obtained from Dropletex (Toronto). Liquid solutions were introduced into the channels using two syringe pumps. Flow speeds were adjusted for the varying solution conditions MCP batches to obtained droplet of about 10 µm diameter. Typical flow speeds were 0.1-1 µL/min for the aqueous MCP phase and 1-10 µL/min for 1-octanol.

**NADH fluorescence measurements**

Droplets were prepared with 1-octanol carrying 1,2 PD with a concentration of 1.31 mM and imaged right after preparation inside a 96 well plate. NADH was excited using the 405 laser line and fluorescence was collected between 417 and 477 nm.

**HPLC measurements**

MCP-containing droplets were produced as described using microfluidic methods. In a 1.5 mL Eppendorf tube, 15 µL MCP emulation was covered by 150 µL 1-octanol supplemented with 4 mM 1,2-PD. Either passive or active MCP were used that were labeled "passive W/O" or "active W/O" in Fig. 3C. Control samples were prepared by 15 µL MCP in aqueous buffer that was brought into contact with a 1-octanol phase in the same way as the emulsion samples and was labeled “active water”. Solution conditions and co-factor concentrations were otherwise as described above. All three samples remained phase separated over the experimental time frame. After four hours incubation at room temperature the 1-octanol phase was gently mixed by pipetting, centrifuged at 10,000 g to fully separate from the aqueous phase and a sample of the 1-octanol phase was harvested and transferred for further analysis using HPLC. Samples were run on an Agilent 1260 HPLC system. Metabolites were separated using a Rezex ROA-Organic Acid H+ (8%) LC column (Phenomenex) with 5 mM sulfuric acid as the mobile phase flowing at 0.4 mL/min at 35 °C. Peak areas were calculated using Agilent ChemLab software.

**Droplet polarization and chemotaxis experiments**

100 µL pre-wetted 1-octanol was pipetted into a 96-well plate. Then a 0.5 µL aqueous droplet containing 0.001 mg/ml Atto 647N and 13.1 mM 1,2 PD was placed into the well, establishing a dye and 1,2-PD containing diffusion front. Into this diffusion front, MCP containing droplets were pipetted. To obtain droplet between 10 and 100 µm in diameter, the plastic tip of the pipette tip was pressed against the glass surface when a volume of 0.1 µL was ejected. Individual droplets were then imaged over time.

**Confocal imaging and image analysis**

Confocal images were obtained on a Nikon Eclipse Ti2 microscope using 20 or 40x lens. The NADH-reactive molecule was excited at 405 nm, GFP at 488 nm, mCherry at 560 nm and Atto 647N at 640 nm using solid-state lasers. Images were quantified using Fiji version 2.1.0/1.53c (Schindelin et al., 2012). For data shown in Fig. 3B a circular region of interest (ROI) of 5 µm diameter centered in each MCP droplet was quantified in NADH intensity over time. The intensity ratio in Fig. 4 C were obtained using the Radial Profile Extended plugin (Philippe Carl). Specifically, the average GFP or mCherry intensity a 45° pie-shaped segment of the circular MCP droplet was calculated. The intensity ratio was then calculated between the segment facing towards and the segment pointing away from the 1,2-PD source (as determined by the Atto 647N fluorescence). Also see sketch insert in Fig. 4C where the segments are shown shaded in gray. Using the same procedure NADH fluorescence ratio was quantified (Fig. 4 D). The example micrographs in panel Fig. 4C were contrast enhanced by applying a Gaussian blur filter, background subtraction and histogram stretching where 0.35% of all pixels were saturated. Image quantification was otherwise performed on raw images.

#### Supplemental Figures

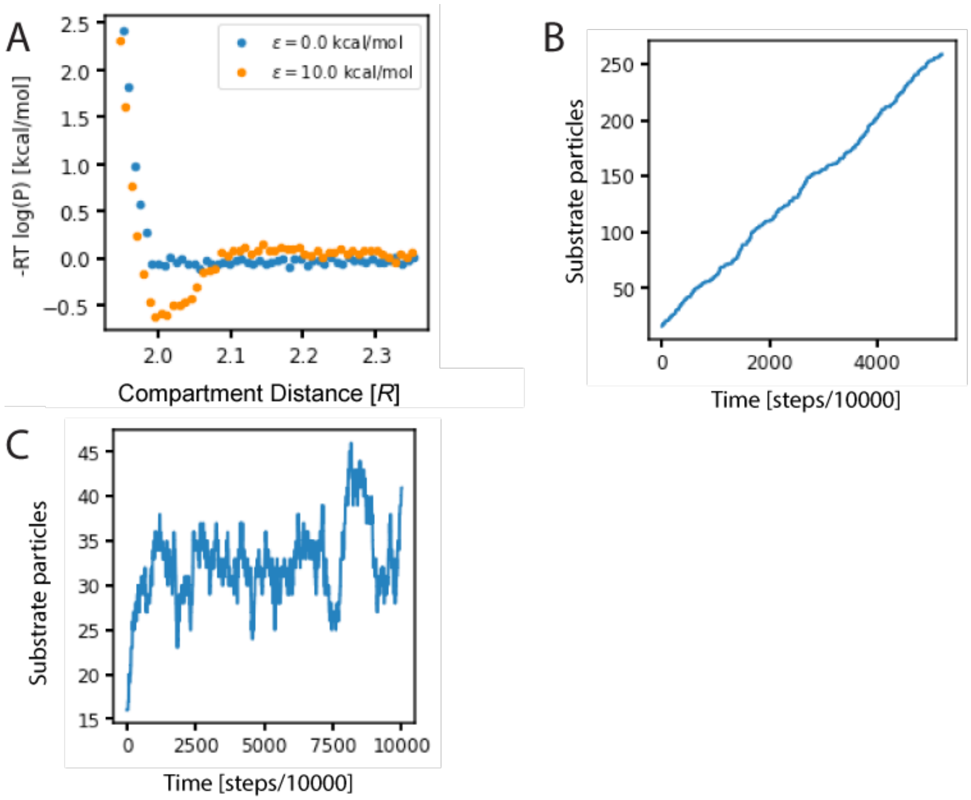


**Fig S1** – A) Compartment-Compartment effective interaction potential in the presence of n=20 particles. B) Example trajectories that show the substrate concentration for passive particles $k_c$=0 $k_s$= 0.005 x $10^{-3}$/steps. C) Active compartments $k_c$ = 0.0001 x $10^{-3}$/steps and $k_s$=0.005 $10^{-3}$/steps.

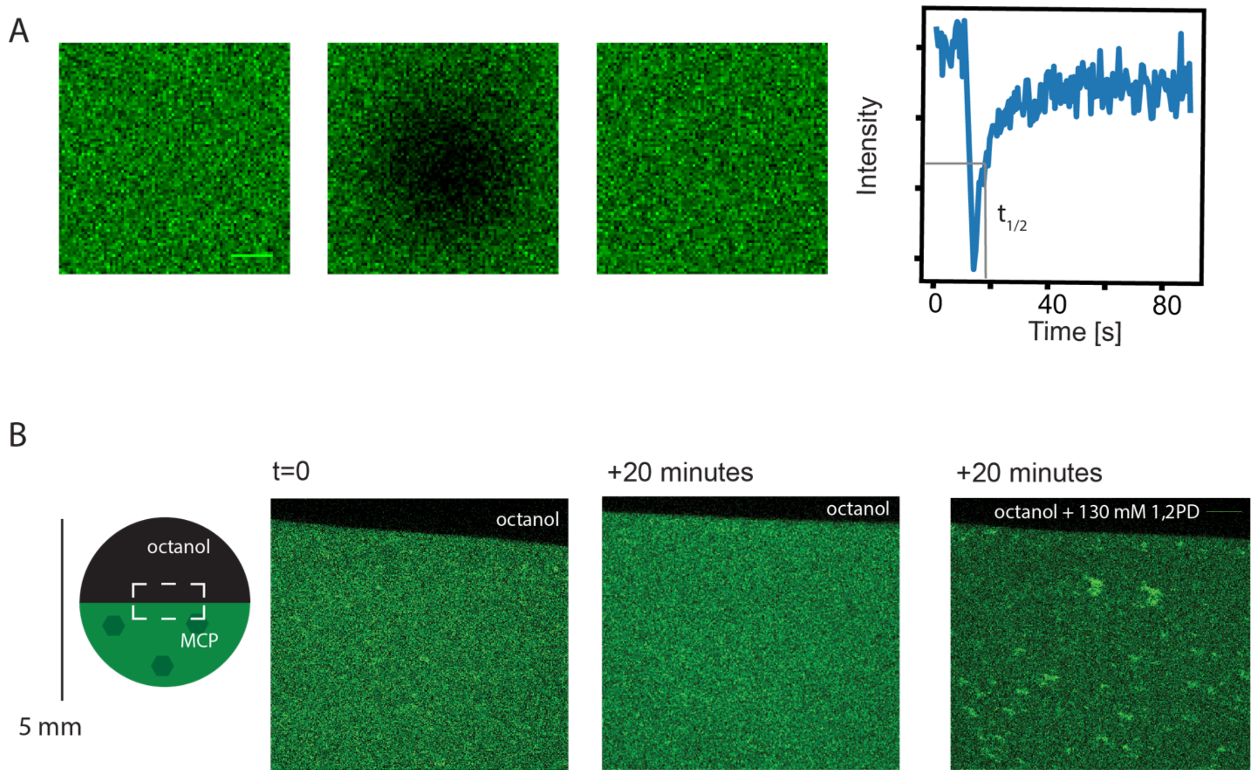


**Fig S2** -A) Example trajectory of FRAP experiment with three representative images before (left), immediately after (middle), and after full recovery (right) from bleaching laser pulse. The half time of recovery is indicated in the figure as $t_{1/2}$. B) Addition of 1,2-PD via diffusion from the octanol phase into the MCP-containing water phase (MCP fluorescent shown in green). MCP aggregates are seen as bright spots after 20 minutes incubation time (rightmost image).

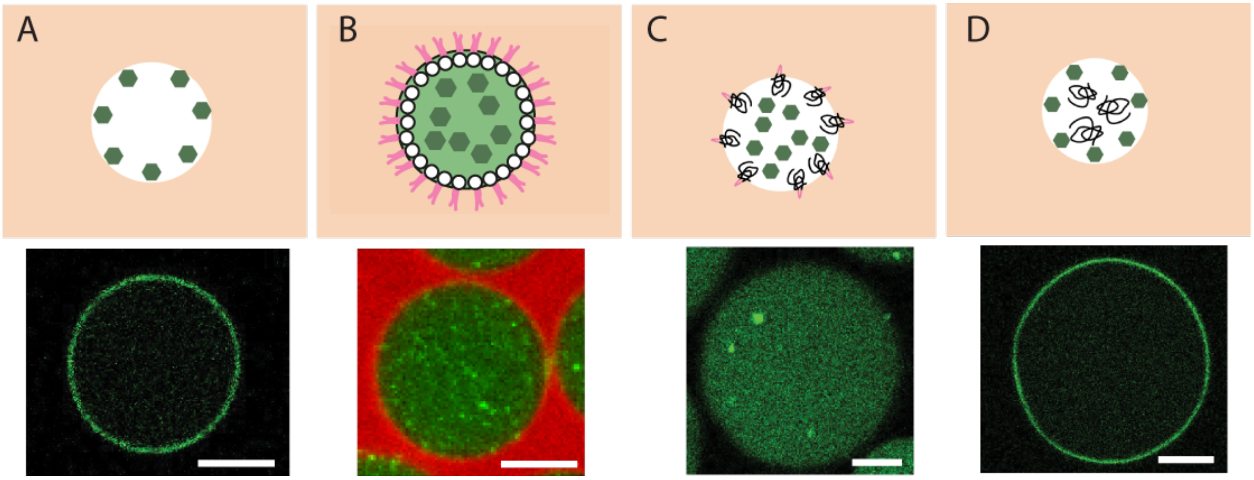


**Fig S3** - MCPs in water droplets (white in all cartoons) and in 1-octanol (light orange in all cartoons) for different water and oil compositions: A) Water phase with phosphate buffered saline (PBS); B) Water phase with PBS and 10 mg/ml dissolved DOPC lipid in 1-octanol; C) Water phase with 100 mg/ml F68 polymer in PBS D) Water phase with 100 mg/ml PEG (MW 8 kDa) polymer in PBS. Bottom panel shows the corresponding confocal images with MCP shown in green and lipids in red. Scale bars 35 µm, 10 µm, 10 µm and 30 µm.

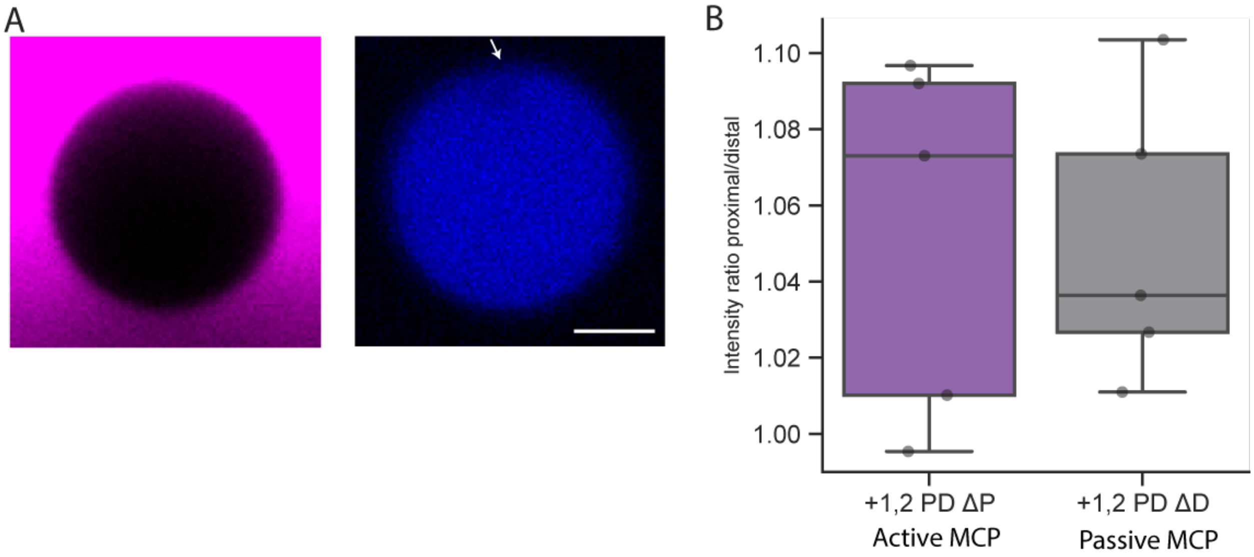


**Fig S4** – A) NADH fluorescence within active compartments. Dye (cyan) is a proxy for the 1,2-PD gradient. Arrow points to a local decrease in NADH concentration, indicating localized activity at the proximal site (with higher MCP concentration). B) Comparison of chemotaxis for active (cyan) and passive MCPs (gray) within the same droplet. Note that both populations indicate chemotaxis with values > 1.0. Scale bar 25 µm.

### Supplementary Theory – Enzymatically-active bacterial microcompartments follow substrate gradients and are protected from aggregation

#### I. MAIN STARTING EQUATIONS

In Ref. 1 the transport properties of monomers that can reversibly dimerize were studied. It was found found that, assuming local binding equilibrium, the concentration of free monomers $\rho_1$ and dimers $\rho_2$ is related to the total, summed up concentration of monomers and dimers $\rho_{\text{tot}} = \rho_1/2 + \rho_2$ by

$$\rho_1 = \frac{K}{4}\left(\sqrt{1 + 16\frac{\rho_{\text{tot}}}{K}} - 1\right) \tag{1}$$

$$\rho_2 = \frac{\rho_1^2}{K} \tag{2}$$

where $K(x)$ is the (potentially position-dependent) binding equilibrium constant between monomers and dimers. In the context of MCP chemotaxis, the binding equilibrium constant decreases with increasing concentration $c_{\text{s}}$ of the substrate 1,2-PD, so that $K(x)$ is a decreasing function of $x$ if $c_{\text{s}}(x)$ is an increasing function of $x$.

Moreover, the effective diffusion coefficient of the total concentration $\rho_{\text{tot}}$ was found to be

$$D_{\text{eff}} = D_2 + \frac{D_1 - D_2}{\sqrt{1 + 16\rho_{\text{tot}}/K}} \tag{3}$$

where $D_1$ and $D_2$ are the diffusion coefficients of the monomers and dimers, respectively.

Lastly, it was found that in steady state the monomer and dimer concentrations must satisfy

$$\frac{1}{2}D_1\rho_1(x) + D_2\rho_2(x) = C \tag{4}$$

where $C$ is a position-independent constant.

#### II. WEAK ASSOCIATION LIMIT (ACTIVE MCPS)

To simplify things further, we consider the case where binding is not very strong ($\rho_{\text{tot}} \ll K$) so that we have mostly monomers and just a few dimers This is justified for active MCPs which self-regulate the local concentration of the substrate that induces the cross-linking. Eqs. (1-2) then simplify to

$$\rho_1 \approx 2\rho_{\text{tot}}\left(1 - 4\frac{\rho_{\text{tot}}}{K}\right) \tag{5}$$

$$\rho_2 \approx 4\frac{\rho_{\text{tot}}^2}{K} \tag{6}$$

Eq. (3) simplifies to

$$D_{\text{eff}} \approx D_1\left(1 - 8\alpha\frac{\rho_{\text{tot}}}{K}\right) \tag{7}$$

with $\alpha \equiv (D_1 - D_2)/D_1$.

Finally, substituting Eqs. (5-6) into Eq. (4) we obtain

$$\rho_{\text{tot}}(x) - 4\alpha\frac{\rho_{\text{tot}}^2(x)}{K(x)} \approx C \tag{8}$$

#### III. PROXIMAL VS DISTAL CONCENTRATIONS IN THE WEAK ASSOCIATION LIMIT

Let us now focus on two points in space, P (proximal) and D (distal), with $c_s(x_P) > c_s(x_D)$ and therefore $K(x_P) < K(x_D)$, so that stabilitaxis will cause $\rho_{tot}(x_P) > \rho_{tot}(x_D)$ in steady state, i.e. accumulation in P relative to D. The steady state condition (8) implies

$$\rho_{tot}(x_P) - 4\alpha \frac{\rho_{tot}^2(x_P)}{K(x_P)} \approx \rho_{tot}(x_D) - 4\alpha \frac{\rho_{tot}^2(x_D)}{K(x_D)} \tag{9}$$

We define for convenience $\rho_{tot}(x_P) \equiv \bar{\rho} + \delta\rho$ and $\rho_{tot}(x_D) \equiv \bar{\rho} - \delta\rho$ and assume that the difference between the two is not very large, i.e. $\delta\rho \ll \bar{\rho}$. Introducing this into (9) and solving for $\delta\rho$, we obtain to lowest order

$$\delta\rho \approx 2\alpha\bar{\rho}^2 \left( \frac{1}{K(x_P)} - \frac{1}{K(x_D)} \right) \tag{10}$$

or equivalently

$$\rho_{tot}(x_P) - \rho_{tot}(x_D) \approx 4\alpha\rho_{tot}^2 \left( \frac{1}{K(x_P)} - \frac{1}{K(x_D)} \right) \tag{11}$$

Note that it does not matter whether we evaluate the $\rho_{tot}^2$ on the right hand side of this equation at $x_P$ or $x_D$, as this only results in a higher order correction.

Finally, we can use (7) to rewrite (11) as

$$\frac{\rho_{tot}(x_P) - \rho_{tot}(x_D)}{\rho_{tot}} \approx \frac{D_{eff}(x_D) - D_{eff}(x_P)}{2D_1} \tag{12}$$

which relates the effective diffusion coefficients to the steady state relative concentrations at each location.

If we assume that at point D the MCPs are almost entirely in the monomer state so that $D_{eff}(x_D) \approx D_1$, we can rewrite (12) as

$$\frac{\rho_{tot}(x_P)}{\rho_{tot}(x_D)} \approx 1 + \frac{1}{2} \left( 1 - \frac{D_{eff}(x_P)}{D_{eff}(x_D)} \right) \tag{13}$$

In particular, for $\frac{D_{eff}(x_P)}{D_{eff}(x_D)} \simeq 0.8$, this gives $\frac{\rho_{tot}(x_P)}{\rho_{tot}(x_D)} \simeq 1.1$, in good agreement with the experimental results in Fig. 4B,C of the main text, without any adjustable parameters.

[1] J. Agudo-Canalejo, P. Illien, and R. Golestanian, Proceedings of the National Academy of Sciences **117**, 11894 (2020).
